# Antidepressant fluoxetine engages astrocytic cAMP via purinergic signalling

**DOI:** 10.64898/2026.06.04.730071

**Authors:** Catriona Marston, Kratika Mujmer, Bárbara Vaccari Cardoso, Sergey Kasparov, Anja G Teschemacher, Valentina Mosienko

## Abstract

The use of selective serotonin reuptake inhibitors (SSRIs), the first-line treatment for depression, has increased by about 50% over the past decade, placing them amongst the top 10 most frequently prescribed drug classes globally. Overall, SSRIs are effective in reducing frequency, severity, and duration of depressive episodes for a majority of patients, yet the mechanisms underlying their therapeutic effects are not fully understood. While SSRIs elevate synaptic serotonin, this action alone cannot account for their therapeutic effects. Additionally, SSRIs engage astrocytes, enhancing cyclic adenosine monophosphate (cAMP) signalling which is reported to be downregulated in depression. However, the signalling mechanisms underlying SSRI-induced upregulation of the astrocytic cAMP pathway remain unclear. Here, we identify a cascade of events by which the SSRI fluoxetine elevates intracellular cAMP levels in astrocytes, a process that depends on astrocyte-microglia crosstalk and purinergic signalling. Using FRET-based sensors in primary rat astrocytes, we show that fluoxetine elevates intracellular cAMP by 28% without altering calcium dynamics. cAMP increase was blocked by both serotonin (5-HT) 2B and adenosine 2B (A2B) receptor antagonists. Using the GRAB-ATP1.0 sensor and luminescence assays, we revealed that fluoxetine enhances astrocytic ATP release by 10% in a 5-HT2B receptor-dependent manner. Consistent with microglia-driven conversion of extracellular ATP to adenosine, which engages astrocytic A2B receptors, depletion of microglia in astrocyte cultures diminished fluoxetine-induced cAMP elevations and increased extracellular ATP. Together, these findings reveal that fluoxetine requires glial crosstalk and coordinated purinergic signalling to enhance astrocytic cAMP, a process shown to contribute to the therapeutic effect of SSRIs.

## Introduction

Selective serotonin reuptake inhibitors (SSRIs) are the first line of treatment for depressive disorders that represent, with ∼5.7% of the population affected, the most common mental health condition and the leading cause of disability worldwide (1). SSRIs are amongst the top 10 most frequently prescribed drug classes (2), with prescriptions increasing by 46% over the past decade (3). SSRIs alleviate depressive symptoms in 40-60% of patients, although evaluating their efficacy is challenging due to the heterogeneity in symptom severity and clinical presentation (4,5). Overall, SSRIs are widely recognised as effective in reducing the frequency, severity, and duration of depressive episodes (6). While some studies have questioned their overall efficacy (7), findings evaluating efficacy are often limited by methodological challenges, including difficulties in measuring treatment response, the absence of reliable biomarkers, and high placebo response rates in clinical trials.

SSRIs increase extracellular concentration of serotonin (5-hydroxytryptamine, 5-HT) by inhibiting its reuptake through the neuronal serotonin transporter (SERT), a mechanism initially proposed to underlie their therapeutic effects (8). Although SSRIs exhibit high affinity for SERT, it is still debated whether this action alone mediates their clinical efficacy or represents just one component of their mechanism of action. Fluoxetine (FLX), one of the most commonly prescribed SSRIs, has high affinity for SERT (K_i_≈5.2×10⁻⁸ M) and substantially lower affinity for other monoamine transporters, including dopamine (≈10^-5^ M) and noradrenaline (≈1.5×10^-5^ M, (9)). However, lack of brain serotonin does not exacerbate depression-like phenotype, nor does it interfere with antidepressive effects of SSRIs in preclinical models of depression (10–12). These findings suggest that although SSRIs increase brain serotonin levels, their therapeutic effect may be mediated via mechanisms independent of 5-HT availability.

One potential alternative pathway may be the activation of 5-HT2B receptors to which FLX has fairly high affinity (5-10×10-6 M), and which were shown to be required for antidepressive properties of SSRIs (13,14).

SSRIs act not only on neurons but also affect astrocytes, a major population of non-neuronal cells in the brain. Evidence from post-mortem human studies and animal models of depression suggests that SSRIs reverse reductions in astrocyte number and associated morphological changes observed in depression, including decreased occupied area and degree of ramification (15–19).

In addition, SSRIs modulate intracellular signalling pathways in astrocytes. Antidepressants have been shown to potentiate astrocytic cyclic adenosine monophosphate (cAMP) signalling by elevating intracellular cAMP levels and promoting the activation of cAMP response element-binding protein (CREB) (20). cAMP is an ubiquitous second messenger in the central nervous system, generated from adenosine triphosphate (ATP) by adenylate cyclase, and both cAMP levels and adenylate cyclase expression have been extensively studied in the context of depression (21). Expression of adenylate cyclase 2, one of ten adenylate cycle isoforms, is decreased in a mouse model of anxiety and also dysregulated in patients with depression (22,23). Some studies suggest that astrocytic cAMP-dependent signalling may contribute to SSRI-induced synaptic plasticity, in part through increased expression of brain-derived neurotrophic factor (BDNF) (24–27). However, the mechanisms by which antidepressants such as SSRIs potentiate cAMP signalling in astrocytes remain incompletely understood. One potential pathway under discussion is the SSRI-induced relocation of Gαs from lipid raft domains to non-raft regions of the plasma membrane, thereby facilitating activation of cAMP signalling pathways (24,28,29).

Another study suggested that antidepressant-induced activation of the cAMP-CREB pathway requires P2Y11 and adenosine 2B (A2B) receptors, and ATP release dependent on the vesicular nucleotide transporter (VNUT) (30). Genetic deletion of VNUT selectively in astrocytes markedly attenuated both fluoxetine-induced ATP release and antidepressant-like behavioural effects, establishing a causal link between astrocytic purinergic signalling and therapeutic efficacy (30). However, FLX targets mediating ATP release from astrocytes and signalling events linking ATP release to cAMP elevation in astrocytes are unknown.

In this study, we show that FLX-driven elevation of cAMP in astrocytes requires microglia-astrocyte crosstalk and is mediated by purinergic signalling. Specifically, we discovered that SSRI FLX engages astrocytic 5-HT2B receptors for ATP release. Extracellular ATP then is converted by microglia to adenosine, which, in turn, acts on astrocytic Gs-coupled A2B receptors, resulting in elevated astrocytic cAMP.

## Methods

### Primary glial cell culture

All procedures complied with the UK Home Office (Scientific Procedures) Act (1986) and were approved by the University of Bristol ethical committee (UIN25050). Primary astrocytes were prepared as described previously (31). Briefly, Wistar Han IGS female and male rat pups at postnatal days 3-8 were culled using schedule 1 method, and midbrain tissue was immediately dissected and dissociated in HBSS (Gibco, 14175129) containing 3 mg/mL BSA (Sigma-Aldrich, A3294), 0.04 mg/mL DNase I (Merck, D5025), and 0.025 mg/mL trypsin (Sigma-Aldrich, T9935). Cells were maintained in T75 flasks in DMEM GlutaMAX (Gibco, 61965059) supplemented with 10% FBS (heat inactivated, Gibco, 12483020) and 1% penicillin-streptomycin (Sigma-Aldrich, P4333) at 37 °C and 5% CO₂.

Primary astrocytes were treated with 10µM of FLX, a dose chosen from human studies showing that at a clinically low dose of FLX (20mg/kg) its concentrations in the brain reaches 10 to 30µM throughout a course of treatment (32) while higher clinically relevant doses result in FLX concentrations up to 50µM in the human brain (33,34). Previously reported and our own preliminary cell culture experiments (Supplementary Table 1) confirmed that this dose does not affect astrocyte survival (35,36).

### Viral transduction and live-cell imaging

Astrocytes were plated on collagen-coated glass coverslips (2 × 10⁵ cells/mL, 24-well plates) and transduced at seeding with adenoviral vectors (AVV) encoding sensors (MOI 50-100): AVV-mCMV/GfaABC1D-Epac-S^H187^ (FRET; cAMP; gift from Kees Jalink and Anne Lann) (37), AVV-mCMV/GfaABC1D-Twitch2B (FRET; Ca²⁺; gift from Oliver Griesbeck) (38), and AVV-CMV-GRAB-ATP1.0 (GRAB-ATP1.0; gift from Yulong Li, Addgene plasmid #167582; http://n2t.net/addgene:167582; RRID:Addgene_167582) (39). Imaging was performed after 2-4 days post-transduction.

Time-lapse imaging was conducted on a Leica SP5 confocal microscope (40× water objective) in a temperature controlled chamber with continuous superfusion of HEPES-buffered saline (pH 7.4) at 2.5ml/min for imaging of FRET sensors or 5ml/min for imaging of GRAB-ATP1.0 sensor. Drugs were delivered by bath. Images were acquired every 2s (GRAB-ATP1.0) or 5 s (FRET) for 1200 s (512 × 512 pixels). FRET sensors were excited by 458 nm Ar laser line; emissions were collected at 465–500 nm (donor) and 515–595 nm (acceptor). Excitation for both the GRAB-ATP1.0 sensor was 488nm, and emissions were collected at 500-600nm. Regions of interest were analysed in LAS X and exported for processing in R. Signals were normalised to baseline of 180s or 300 s and specified in figure legends.

### Determination of ATP release

ATP release was quantified using the RealTime-Glo Extracellular ATP Assay (Promega, GA5010). Astrocytes (6 × 10⁴ cells/well) were plated in opaque 96-well plates, equilibrated in HBSS with 7 mM glucose and treatments, and luminescence was recorded kinetically at 37 °C using a Tecan plate reader (TECAN, infiniteM200pro). Background was subtracted, and values normalised to controls.

### Depletion of microglia

Microglia were depleted using the CSF1R inhibitor PLX5622 (10 µM, 7-10 days; medium refreshed every 3 days). Cultures were maintained in PLX-supplemented media prior to experiments, and depletion was verified by loss of microglial marker ionised calcium-binding adaptor molecule 1 (Iba-1).

### Immunohistochemistry

Coverslips seeded with glia culture 24 hours prior (2×10^5^ cells/ml) were fixed with 4% paraformaldehyde solution (15 min, room temperature (RT)) and then washed with PBS. Cells were permeabilised in blocking solution (10% FBS and 0.3% Triton-X100 in PBS; 30 mins, RT) and then incubated with primary antibodies against astrocyte-specific glial fibrillary acidic protein (GFAP) (anti-GFAP, rabbit, Dako, GA524, 1:500) or microglia-specific Iba1 (anti-Iba1, guinea pig, Synaptic Systems/234 308, 1:400) in an antibody solution (3% FBS and 0.1% Triton-X100 in PBS) at 4°C for 16 hours. After PBS washes to remove excess primary antibodies, cells were incubated with secondary antibodies, goat anti-rabbit Alexa Fluor 488 (Invitrogen, R37116, 2 drops/ml) or goat anti-guinea pig Alexa Fluor 594 (Invitrogen, A11076, 1:500). Coverslips were air-dried and mounted on slides using FluorSave mounting media (Millipore, 345789). Stained coverslips were imaged using a 20x air objective on a Leica SP5 confocal microscope.

### Statistical analysis

For analysis of data generated using FRET sensors, the FRET ratio was calculated and normalised to the baseline of each recording across all presented data, except in Figure 4D, where non-normalised values are reported. Normalised FRET ratio were calculated using a custom-made R script (V4.1.1). Next, each recording was carefully assessed to ensure that the baseline was stable, and focus was consistent throughout the recording. For both FRET sensor and GRAB-ATP1.0 recordings, the area under the curve (AUC) was calculated from 300 to 1200 seconds using GraphPad Prism (V11).

For all experiments, to determine if the drug(s) tested changed the tested parameter, a one-sample t-test from zero was used. To assess the significance between experimental groups, an unpaired t-test was used for two groups and a one-way analysis of variance (ANOVA) with, if necessary, a post-hoc Holm-Šídák test was used for comparing more than two groups.

## Results

### FLX elevates intracellular cAMP levels without affecting Ca^2+^ dynamics in primary rat astrocytes

Intracellular levels of Ca^2+^ and cAMP following FLX application (10µM, 5 minutes) were quantified using live-cell confocal imaging of Twitch2B and Epac-S^H187^ FRET sensors, respectively, in rat primary astrocytes (Figure 1A&B). FLX had no effect on intracellular Ca^2+^ levels (t(22)=1.487, p=0.1511, Figure 1B&C), in contrast to ATP (10µM, 5 minutes) which triggered a robust elevation by 160% compared to baseline (t(19)=8.311, p<0.0001), verifying that the sensor accurately reports Ca²⁺ changes (Figure 1B&C).

**Figure 1:**
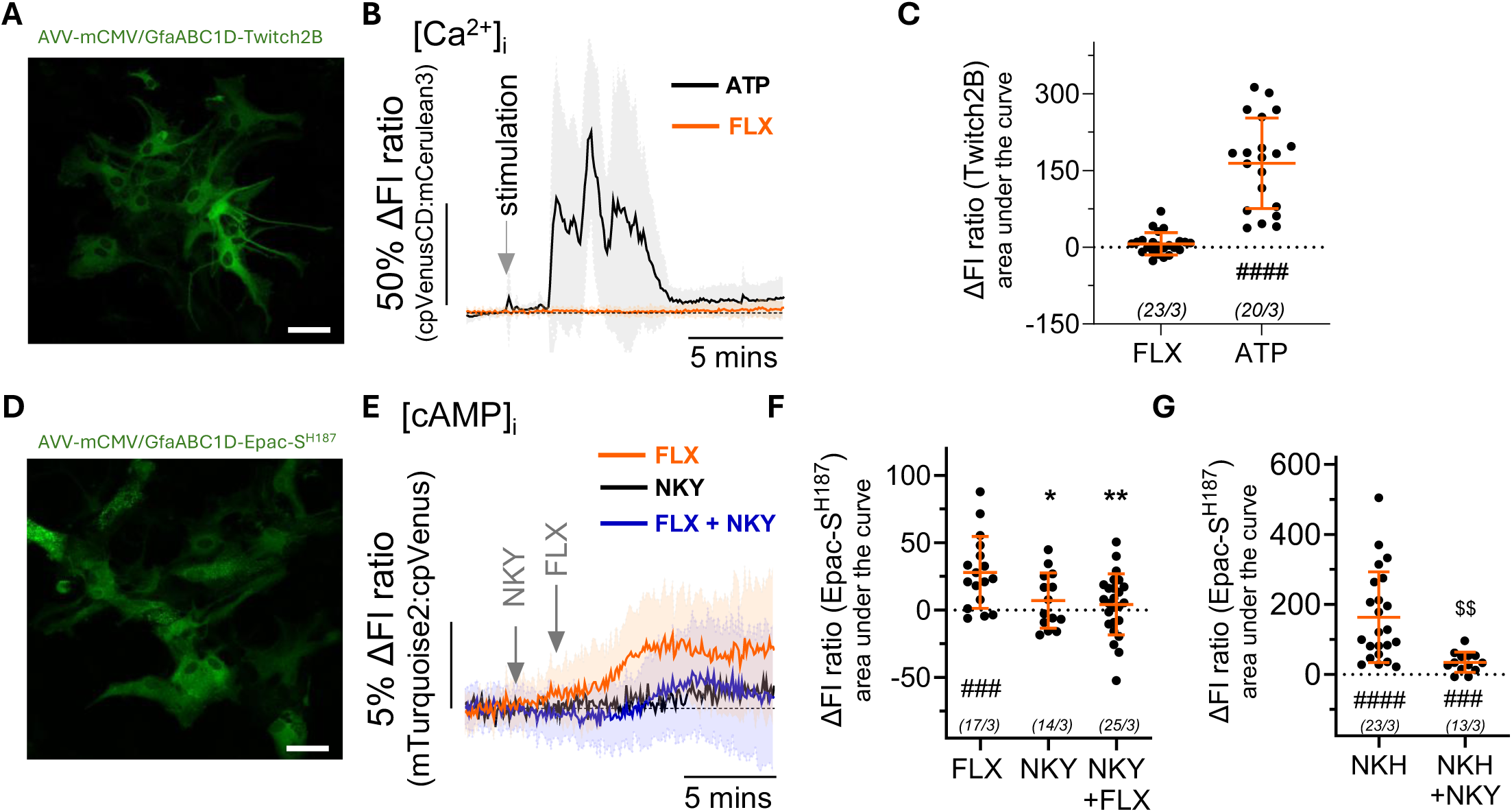
Fluoxetine enhances astrocytic cAMP signalling. **(A)** Representative image of primary rat astrocytes transduced with adenoviral vectors (AVVs) to express intracellular calcium FRET sensor Twitch2B (AVV-mCMV/GfaABC1D-Twitch2B). Average traces **(B)** and summary data **(C)** illustrating changes in intracellular calcium levels ([Ca^2+^]_i_; Twitch2B; fluorescence intensity (FI) ratio of cpVenus:mCeruleuan3) in primary rat astrocytes following application of fluoxetine (FLX, 10µM, 5mins) or ATP (1µM, 5mins). **(D)** Representative image of primary rat astrocytes transduced with AVVs to express intracellular cAMP sensor Epac-S^H187^ (AVV-sGFAP-cAMPhigh^H187^). Average traces **(E)** and summary data **(F)** illustrating changes in intracellular cAMP levels ([cAMP]_i_; Epac-S^H187^; FI ratio of mTurquoise2:Venus) in primary rat astrocytes following application of FLX (10µM, 5mins) along or in combination with adenylate cyclase inhibitor NKY80 (NKY, 10µM; NKY 180-1200s and FLX 300-480s), or NKY alone (10µM; 180-1200s). **(G)** Summary data illustrating changes to [cAMP]_i_ following application of adenylate cyclase activator NKH477 (NKH, 3µM, 300-480s) alone or in combination with adenylate cyclase inhibitor NKY80 (NKY, 10µM; NKY 180-1200s and NKH 300-480s). Scale bar = 50µm. Data are presented as mean ± standard deviation with each data point representing a region of interest (ROI). The numbers in parentheses indicate the number of individual ROIs and separate cultures (animals). ^####^p<0.0001, ^###^p<0.001 vs zero, one sample t-test; **p<0.01, *p<0.05 vs FLX, ^$$^p<0.01 vs NKH; unpaired t-test (C and G) or one-way ANOVA with post-hoc Holm-Šídák’s test (F).

FLX application elevated astrocytic cAMP levels by 28% compared to baseline (mean AUC: 27.98±26.70; t(16)=4.320, p=0.0005, Figure 1E&F).

Co-application of FLX with the AC inhibitor NKY80 (NKY; 10µM; 17 minutes) blocked FLX-evoked cAMP increase (NKY+FLX vs zero: (t(24)=0.9299, p=0.3617, or vs. FLX: p=0.0044, Figure 1E&F). In comparison, the AC activator, NKH477 (NKH; 3µM, 3 minutes) increased intracellular cAMP in astrocytes by 164% (mean AUC: 163.5±129.2; t(22)=6.067, p<0.0001, Figure 1G), which was attenuated by NKY80 (NKY vs zero: (t(12)=4.361, p=0.009, vs. NKH: t(34)=3.518, p=0.0013, Figure 1G).

### FLX-driven increase in astrocytic cAMP requires both the A2B and 5-HT2B receptors

To delineate the signalling mechanisms underlying the FLX-induced increase in intracellular cAMP in astrocytes, we examined the potential involvement of receptors previously implicated in the regulation of astrocytic cAMP levels or to which FLX was reported to have affinity.

First, we tested A2B receptor involvement in FLX-triggered cAMP signalling in astrocytes. Functional expression of the A2B receptor has been demonstrated in astrocytes both *in vivo* and *in vitro* in previous studies (40,41), and was also confirmed in our astrocyte cultures (Supplementary Table 2). As the A2B receptor is Gs-coupled, it would be expected to lead to an increase in intracellular cAMP levels if FLX were to engage it.

FLX-triggered increase in astrocytic cAMP was blocked by a highly selective antagonist of A2B receptor (PSB603 (PSB); 10µM; FLX+PSB vs. zero: t(28)=0.4130, p=0.6828; vs. FLX: p<0.0001; Figure 2A&B). We confirmed that PSB is able to block A2B-driven cAMP elevation (PSB+adenosine vs. zero: t(17)=0.6917, p=0.4985; vs. ADO: p=0.0029;) as a response to its intrinsic ligand adenosine (by 113%; adenosine vs. zero: t(49)=8.889, p<0.000110µM; Figure 2D).

**Figure 2:**
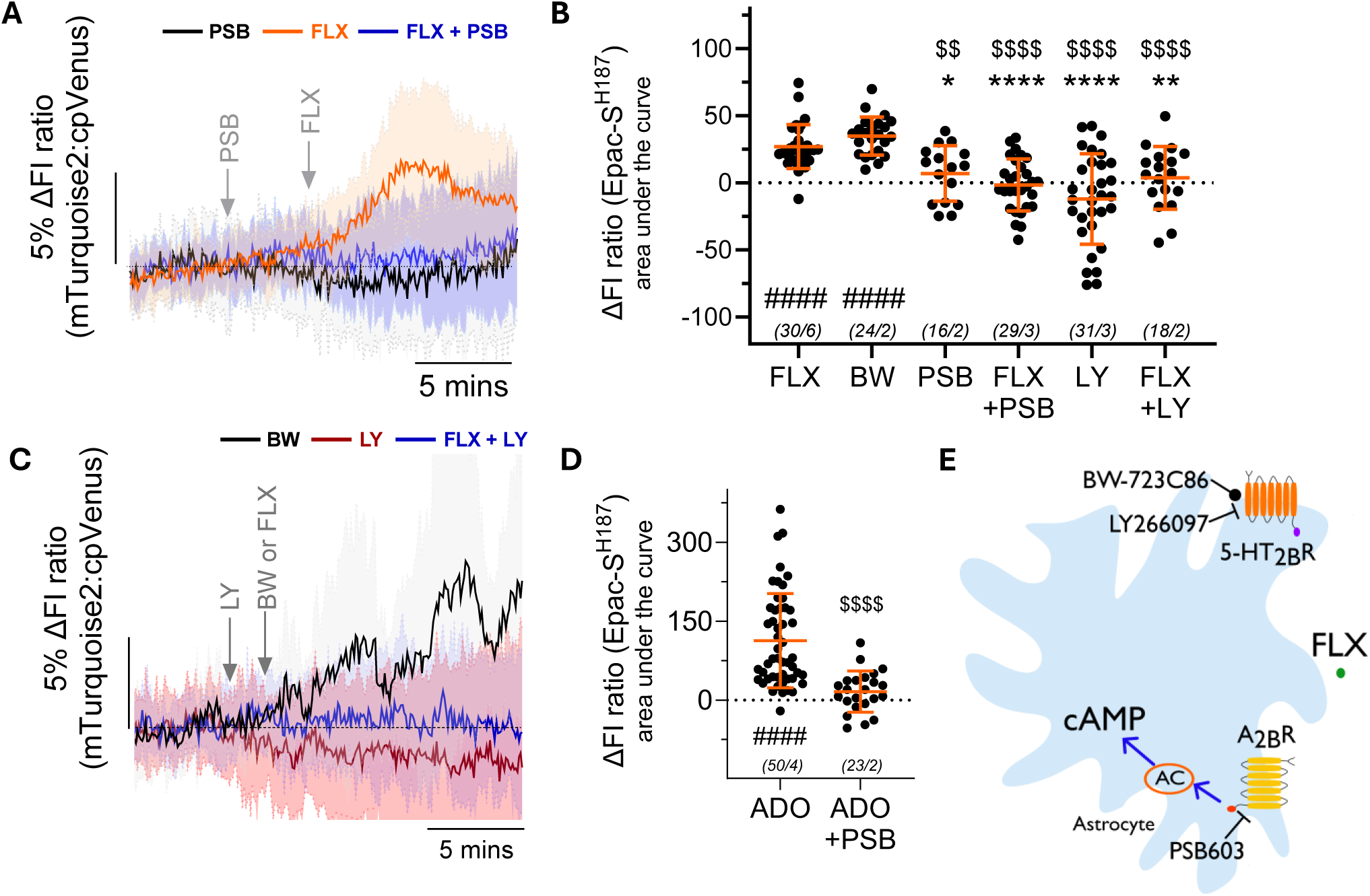
The fluoxetine-triggered increase in astrocytic cAMP requires both adenosine A2B and serotonin 5-HT2B receptors. Average traces **(A)** illustrating changes in intracellular cAMP levels (Epac-SH187; fluorescence intensity (FI) ratio) in primary rat astrocytes following application of FLX (10µM, 5mins), or in combination with adenosine 2B (A2B) receptor antagonist PSB603 (PSB, 10µM, 300-1200s). Summary data **(B)** illustrating changes to [cAMP]_i_ following application of FLX, BW, PSB, FLX+PSB, LY, and LY+FLX. Average traces **(C)** illustrating changes to [cAMP]_i_ following application of FLX alone, or in combination with serotonin 2B receptor (5-HT_2B_R) antagonist LY266097 (LY, 10uM, 300-1200s) or 5-HT_2B_R agonist BW723C86 (BW,1µM, 5mins). **(D)** Summary data illustrating changes to [cAMP]_i_ following application of adenosine (ADO, 10µM, 480-600s) alone or in combination with A_2B_R antagonist (PSB, 10µM, 300-1200s). **(E)** Schematic illustration of signalling pathways involved in FLX-driven recruitment of astrocytic cAMP: AC – adenylate cyclase; round arrow – agonist; flat arrow – antagonist. Data are presented as mean ± standard deviation with each data point representing a region of interest (ROI). The numbers in parentheses indicate the number of individual ROIs and separate cultures (animals). ^####^p<0.0001 vs zero, one sample t-test; ****p<0.0001, **p<0.001, *p<0.01 vs FLX, ^$$$$^p<0.0001, ^$$^p<0.001 vs BW (C), one-way ANOVA with post-hoc Holm-Šídák’s test; ^$$$$^p<0.0001 vs ADO (D), unpaired t-test (D).

Notably, the temporal dynamics of cAMP responses differ between adenosine and FLX. FLX requires significantly longer time to reach the maximal response compared to adenosine (mean time (s) ± SD: ADO = 657.2±129.4s; FLX = 906.2±340.9s; t(31)=2.739, p=0.0101), indicating that FLX-stimulated increase in cAMP is indirect, and suggesting that FLX engages another target upstream of the A2B receptor. We therefore tested whether the 5-HT2B receptor, to which FLX has high affinity, could contribute to the fluoxetine-induced cAMP elevation in astrocytes. This is feasible since 5-HT2B receptor expression has been found in murine astrocytic cultures (42) and human astrocytes (43), and in our rat astrocyte cultures (Supplementary Table 2 and 3; Supplementary Figure 1).

FLX-induced increase in astrocytic cAMP was abolished by a specific 5-HT2B receptor antagonist LY266097 (LY; 10µM; LY vs. zero: t(17)=0.6917, p=0.4985; vs. FLX: p=0.0029, Figure 2B&C). While application of 5-HT2B receptor agonist BW-723C86 (BW; 1µM; 5 minutes) significantly elevated intracellular astrocytic cAMP by 40% (t(22)=11.760, p<0.0001) and to an extent comparable to that induced by FLX (t(37)=2.007, p=0.0521; Figure 2B&C).

Altogether these data indicate that FLX-induced elevation of cAMP in astrocytes requires coordinated engagement of two receptor systems, with A2B receptor activation occurring downstream of 5-HT2B receptor signalling (Figure 2E). We hypothesised that stimulation of the 5-HT2B receptor results in autocrine activation of A2B receptors by adenosine. It is well established that a significant proportion of extracellular adenosine comes from conversion of ATP by extracellular nucleases (44). Therefore, we next investigated whether FLX can evoke ATP release from astrocytes.

### FLX engages 5-HT2B receptors to trigger ATP release in primary rat astrocytes

To quantify extracellular ATP dynamics following FLX treatment, we first employed the genetically encoded fluorescent sensor GRAB-ATP1.0. FLX application produced a significant increase in extracellular ATP levels by 10% (t(28)=2.111, p=0.0438; Figure 3A). However, given the rapid kinetics and spatial complexity of ATP signalling, as well as the continuous basal release of ATP by astrocytes, which can further propagate ATP release in neighbouring cells (43), it was challenging to reliably quantify ATP dynamics following pharmacological treatments using GRAB-ATP1.0. We therefore complemented these experiments with a real-time ATP luminescence assay to independently validate the findings. Consistent with the extracellular sensor data, FLX treatment significantly increased ATP release relative to control by 2% (FLX vs. control p=0.0082; Figure 3B&C). Given complexity of astrocyte ATP signalling combined with the large assay volume (100µl), accurately measuring short-term local ATP dynamics is technically challenging using luminescent detection. Although we managed to achieve reliable and reproducible results on ATP release using luminescent assay, which allowed us to test various pharmacological agents, the observed responses were modest and likely do not reflect local intercellular ATP fluctuations.

**Figure 3:**
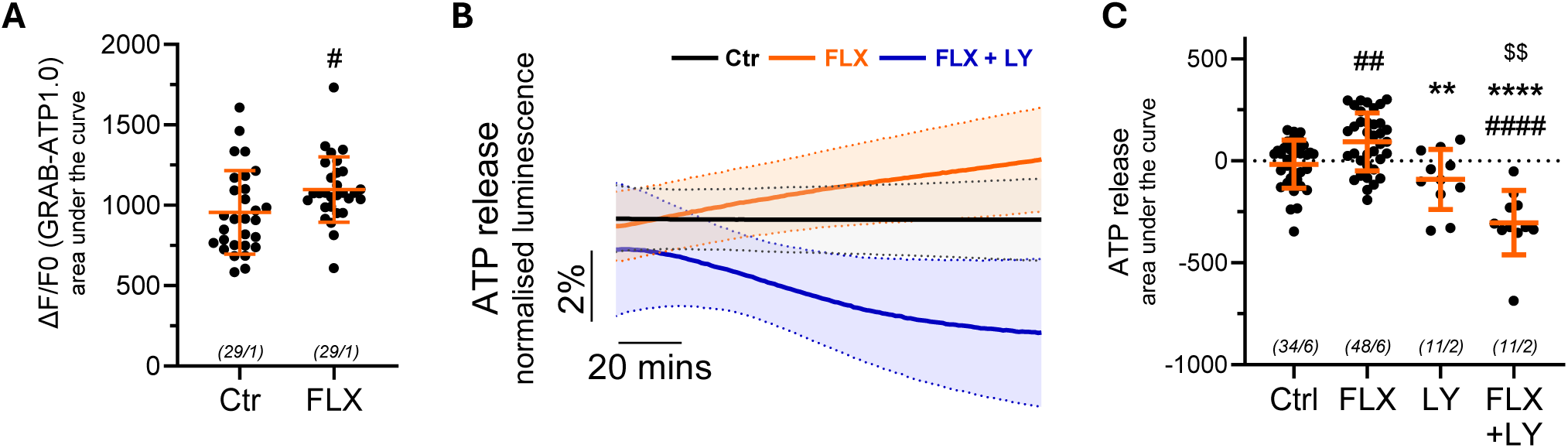
Fluoxetine potentiates astrocytic ATP release via serotonin 2B receptor. **(A)** Amount of ATP released by primary rat astrocytes transduced with AVVs to express extracellular GRAB-ATP1.0 sensor (AVV-CMV-GRAB-ATP1.0) following application of fluoxetine (FLX, 10µM, 300-600s) or HBS as control (Ctr). Average traces **(B)** and summary data **(C)** illustrating changes in extracellular ATP levels following application of FLX (10µM, 4 hours) along or serotonin 2B receptor antagonist LY266097 (LY, 10µM, 5 hrs) or combination of both (LY for 1hr then LY+FLX for 4hrs). Data are presented as mean ± standard deviation with each data point representing a region of interest (ROI; A) or a well (B and C). The numbers in parentheses indicate the number of individual ROIs (A) or well (B and C) and separate cultures (animals). ^####^p<0.0001, ^##^p<0.01, ^#^p<0.05 vs control, ****p<0.0001, **p<0.01, *p<0.05 vs FLX, ^$$^p<0.01 vs LY, paired t-test (A) or one-way ANOVA with post-hoc Holm-Šídák’s test (C).

Inhibition of 5-HT2B receptors with the selective antagonist LY effectively blocked the FLX-induced increase in ATP release. (LY+FLX vs FLX: p<0.0001; Figure 3B&C), demonstrating that this effect is dependent on 5-HT2B receptor activation. Collectively, these data support a model in which FLX engages 5-HT2BR to stimulate ATP release, which in turn acts in an autocrine manner to drive A2B receptor-dependent cAMP production in astrocytes. However, because ATP does not directly bind to the A2B receptor, we next examined whether extracellular ATP is metabolized to adenosine, which could subsequently activate A2B receptor signalling.

### FLX increases astrocytic cAMP by engaging microglia and purinergic signalling

Our rat primary astrocyte cultures were prepared using a standard protocol to minimise microglial contamination (31). However, a residual amount of microglia is still detected in our primary astrocyte preparation (Figure 4A). Microglia play an important role in converting extracellular ATP to adenosine since they are the main cellular source of CD39, which catalyses the first step in that process (45) (Figure 4F). To assess the contribution of microglia to FLX-driven signalling, we treated astrocyte cultures with a microglia-specific CSF1R inhibitor PLX5622 to ablate microglia (45). Indeed, 7-day treatment with PLX5622 (10µM) resulted in near-complete microglial depletion with only an occasional microglial cell detected in the whole field of view (PLX5622-treated vs untreated primary rat astrocyte cultures, p<0.0001; Figure 4A&B).

**Figure 4:**
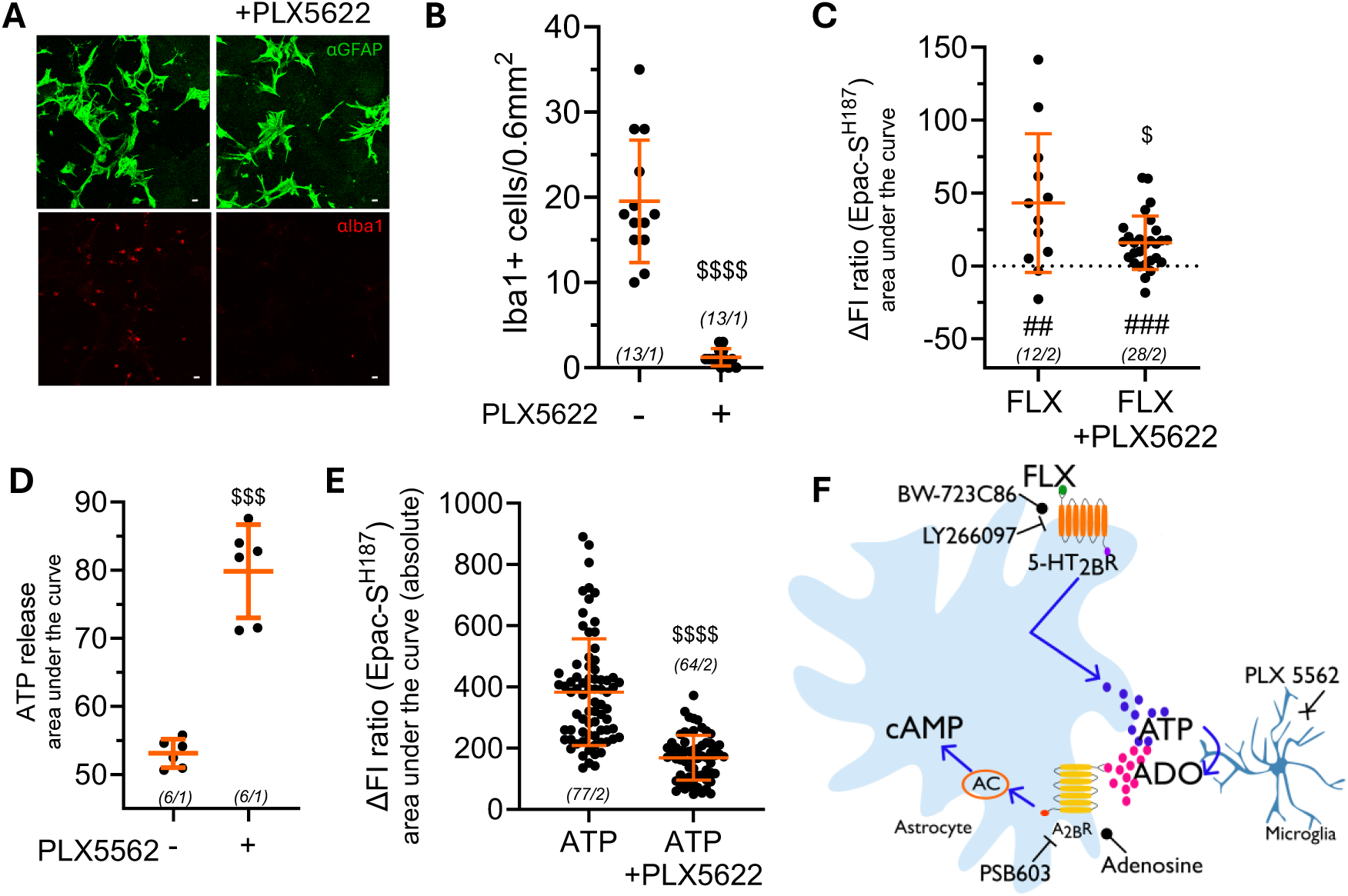
Fluoxetine-driven increase in astrocytic cAMP signaling requires microglia-astrocyte crosstalk mediated by purinergic signaling. Successful microglial depletion by an inhibitor of microglia-specific colony-stimulating factor 1 receptor (PLX5622, PLX, 10µM, 7 days): **(A)** Representative images of staining against astrocyte-specific glial fibrillary acidic protein (GFAP) and microglia-specific ionised calcium-binding adapter molecule 1 (Iba1) and **(B)** quantification of Iba1-positive cells in culture. Scale bar = 20µm. **(C)** Summary data illustrating changes in cAMP levels (Epac-S^H187^; fluorescence intensity (FI) ratio) in primary rat astrocytes with or without PLX treatment following application of fluoxetine (FLX 10µM, 5mins). **(D)** Summary data illustrating changes in extracellular ATP levels in rat astrocyte cultures treated with PLX. **(E)** Summary data illustrating changes in cAMP levels (Epac-S^H187^; fluorescence intensity (FI) ratio) in primary rat astrocytes with or without PLX treatment following application of adenosine triphosphate (ATP, 50µM, 3mins). **(F)** Schematic illustration of signalling pathways involved in FLX-driven recruitment of astrocytic cAMP: Pointed arrow - potentiated; flat arrow - antagonist; rounded arrow - agonist; cross arrow - inhibition; A_2B_R: adenosine 2B receptor; 5-HT2BR – serotonin 2B receptor; AC: adenylate cyclase. Data are presented as mean ± standard deviation, with each data point representing a field of view (B) or region of interest (C and D) (ROI) or well (E). The numbers in parentheses indicate the number of individual field of view (b) or ROI (C and D) or well (E), and separate cultures (animals). ^$$$$^p<0.0001, ^$$$^p<0.001, ^$^p<0.05 vs PLX-free culture, Mann-Whitney test (B), or unpaired t-test (C-E), ^###^<0.001, ^##^<0.01 one-sample t-test from 0 (C).

FLX-induced elevations in astrocytic cAMP were reduced by 62% following microglial depletion (t(38)=2.645, p=0.0118, Figure 4C). However, microglia depletion did not completely abolish FLX-triggered increase in astrocytic cAMP, as there still was a significant increase by 15% in cAMP levels in PLX5622-treated cultures in response to FLX (p<0.0001, Figure 4C).

If microglia substantially contribute to the conversion of extracellular ATP to adenosine, microglial ablation should lead to an accumulation of ATP in response to FLX treatment. Indeed, in PLX5622-treated primary astrocyte cultures, ATP release is 60% higher than in untreated cultures (t(5.906)=9.149, p=0.001, Figure 4D).

Similarly, if microglia plays a major role in ATP-to-adenosine conversion within our experimental system, exogenous ATP conversion into adenosine should be suppressed by microglial ablation, thereby diminishing A2B receptor-dependent cAMP elevation. Indeed, ATP-triggered increase in astrocyte cAMP is reduced by 55% in microglia-depleted cultures compared to astrocytes from regular cultures (t(101.9)=9.605, p<0.0001, Figure 4E).

Together, these findings indicate that microglia plays a critical intermediary role in the ATP-to-adenosine axis underlying FLX-induced cAMP signalling. Specifically, they support a model in which FLX stimulates ATP release from astrocytes, which is subsequently metabolised with the aid of microglia into adenosine, thereby amplifying A2B receptor-dependent cAMP production.

## Discussion

Accumulating evidence highlights astrocytic signalling as a critical component of the antidepressant actions of monoaminergic drugs. SSRIs have been shown to activate astrocytic cAMP signalling, a pathway typically reduced in depression and restored by antidepressant treatment, thereby contributing to their therapeutic efficacy. Yet, the mechanisms underlying antidepressant-induced modulation of astrocytic cAMP remain insufficiently understood. Here, we identify a novel mechanism driving astrocytic cAMP levels following FLX treatment which involves astrocyte-to-microglia crosstalk and purinergic signalling, providing a mechanistic framework for understanding how antidepressants regulate astrocyte function.

Specifically, we show that FLX promotes ATP release in a process dependent on astrocytic 5-HT2B receptors. Microglia, being the main cellular source of CD39, subsequently contribute to the conversion of FLX-induced extracellular ATP into adenosine, which then acts on astrocytic A2B receptors, increasing in intracellular cAMP levels (Figure 4F).

5-HT2B receptors are required for the observed increase in astrocytic cAMP levels. However, we did not detect a concomitant rise in intracellular Ca²⁺, despite their canonical classification as Gq-coupled GPCRs (46). However, the literature indicates that their signalling repertoire is considerably broader and highly context dependent. In addition to classical PLC/IP3/Ca²⁺ signalling, 5-HT2B receptors have been shown to engage ERK/MAPK, PI3K/Akt, β-arrestin-dependent, and receptor-transactivation pathways in a cell type- and ligand-specific manner (47–50). 5-HT2B receptor agonist, BW723C86, has been reported to be unable to evoke Ca^2+^ levels in brainstem astrocytes (51). Consistent with this signalling plasticity, we found that 5-HT2B receptor engagement was required for the fluoxetine-induced increase in astrocytic cAMP despite the absence of a detectable Ca²⁺ rise, indicating that, in our preparation, 5-HT2B receptors are functionally coupled to a non-canonical signalling pathway. Activation of 5-HT2B receptors with the selective agonist BW723C86 produced a significant although indirect response on intracellular cAMP, further demonstrating that 5-HT2B is functional in our preparation and its signalling extends beyond classical Gq coupling. Together, these data establish that fluoxetine increases astrocytic cAMP via a Ca²⁺-independent mechanism, indicating that Ca²⁺ mobilisation is not required for this component of antidepressant action.

The 5-HT2B receptor is expressed by astrocytes throughout the mouse brain. Comparing expression between the striatum, hippocampus and cortex, 5-HT2B receptor is expressed the highest in the striatum, with the lowest expression in the cortex. Relative to GFAP, 5-HT2B receptor expression is around 7-fold lower in the striatum and 9-fold less in the hippocampus (40). In humans, according to the Human Protein Atlas, microglial 5-HT2B receptor expression is double that of astrocytes (52,53). However, in cultures, 5-HT2B receptor expression is higher in astrocytes than in both microglia and neurons (54).

Here, we show that FLX requires 5-HT2B receptor to stimulate ATP release from astrocyte. SSRIs were previously reported to facilitate ATP release specifically from astrocytes via VNUT (30,55) or gap junctions (56). Preclinically, interfering with mechanisms of ATP release induces a depressive-like phenotype that can be reversed by ATP administration, and maintaining ATP levels in the mPFC are also crucial for regulating depressive-like behaviour (57,58). Hertz et al., (2015), demonstrated that FLX engages 5-HT2B receptor to regulate intracellular pathways in astrocytes (59), while other published studies revealed lack of 5-HT2B receptor compromises therapeutic effects of FLX (13,14).

Ectonucleotidases CD39 and CD73 represent the principal enzymatic pathway mediating the extracellular conversion of ATP to adenosine (44). Both enzymes exist in membrane-bound forms and as soluble or extracellular vesicle-associated variants within the extracellular milieu (60,61). Expression patterns are cell-type specific, with CD73 most abundantly expressed in neurons and relatively low levels observed in astrocytes, whereas CD39 is markedly enriched in microglia, with expression reported to be 20 times higher than in astrocytes (45). Therefore, particularly in glial cultures, microglia may play a predominant role in adenosine generation from ATP. Although astrocytes predominate in our cultures, low levels of Iba1 are still detectable (Figure 4; Supplementary Table 2).

Indeed, here we showed that in PLX5622-treated cultures, there is a significant reduction in FLX-and ATP-driven increases in cAMP. While unstimulated ATP release was elevated in PLX5622-treated cultures compared with controls, consistent with impaired ATP catabolism in the absence of microglia and supporting a critical role for microglia in converting extracellular ATP to adenosine.

Microglia contribution to FLX-driven cAMP elevation in astrocytes could go beyond ATP-to-adenosine conversion. It is possible that the 5-HT2B receptor, expressed in both cell types, potentiates ATP release from both astrocytes and microglia following FLX treatment. Furthermore, it has been demonstrated that astrocytic ATP release stimulates microglial ATP release (55).

Expression of the A2B receptor in murine astrocytes varies across brain regions, with the highest expression found in the hippocampus. Relative to GFAP, A2B receptor expression levels are about 4-fold lower in the striatum, 5-fold lower in the hippocampus and 1-fold lower in the cortex (40). In humans, A2B receptor expression is 25 times higher in astrocytes than in both neurons and microglia (52,53). Our data demonstrate that cAMP elevations are a consequence of activation of the A2B receptors following potentiated release of ATP as a result of 5-HT2B receptor activation. Although microglial depletion attenuated the FLX-induced cAMP increase, it did not completely abolish it, suggesting a minor astrocytic contribution to ATP-to-adenosine conversion. Another possibility is that FLX directly promotes adenosine release from astrocytes, a phenomenon observed in astrocytes following ketamine treatment (62).

Our data indicate a mechanism by which alterations to cAMP signalling in depression can be restored. Antidepressants have been reported to potentiate astrocytic cAMP signalling by increasing intracellular cAMP levels and activating CREB (20). Further, expression of adenylate cyclase 2 is decreased in both animal models of anxiety and patients with depression (22,23). Enhanced astrocytic cAMP signalling may also contribute to antidepressant-induced synaptic plasticity, in part by increasing BDNF expression (24–27). Our data provides a mechanism by which FLX restores cAMP levels in astrocytes, identifying a pathway by which antidepressants may recover depressive symptoms. Given the central role of cAMP signalling in regulating astrocyte function and morphology, elevations in cAMP may represent a mechanism through which antidepressant treatment restores astrocytic structure and homeostasis (15,17,19).

We tested a single FLX concentration of 10 µM. This dose was selected because it falls within the range of fluoxetine concentrations reported in the human brain during therapeutic treatment, estimated across studies at approximately 0.3-42 µM (32,63,64). Depending on clinical dose, brain concentrations range from 10-30µM following a treatment of a clinically low dose (20mg/kg), and up to 50µM following higher, clinically relevant doses (120mg/kg) (32–34). Thus, 10 µM represents a physiologically relevant concentration that is also commonly used in experimental models. In addition, previous studies and our own preliminary cell culture experiments indicated that this concentration does not compromise astrocyte viability (35,36). Finally, the reported Ki of the 5-HT2B receptor for FLX is greater than 10 µM, although this remains an area of an ongoing debate (59,65,66).

There is a large volume of literature surrounding the mechanistic properties of antidepressants and the pathophysiology of depression (8). In this study, we demonstrate a mechanism by which FLX alters the signalling dynamics of glia, principally astrocytes. It is clear that the mechanisms of action of antidepressants are complex; in this study, we do not discount any other mechanisms of action of depression, and propose that antidepressants act on multiple pathways simultaneously. Here, we not only identify a pathway through which FLX modulates glial signalling, but also provide support for several proposed mechanisms underlying antidepressant action. First, our findings confirm the involvement of the 5-HT2B receptor, which has been suggested to be an important mediator of antidepressant effects (13,14,42). Second, the participation of microglia is consistent with growing evidence implicating these cells in both the pathophysiology of depression and the mechanisms of antidepressant response (67–70). Our data support previous reports that FLX potentiates ATP release, further aligning with evidence that ATP itself may exert antidepressant-like effects (30,56–58). Finally, our data support the evidence that cAMP signalling is implicated in depression, and the mechanism of action of antidepressants (20,22,23).

While our data focuses on acute signalling events, prior work indicates that chronic antidepressant administration modulates glial purinergic pathways and downstream effectors, including adenosine receptor-linked cAMP pathways, which may contribute to longer-term neuroplastic and behavioural effects of SSRIs beyond immediate second-messenger changes (30).

In summary, our findings identify a previously unrecognised glial mechanism through which fluoxetine modulates astrocyte signalling. We show that FLX engages the 5-HT2B receptor to promote ATP release, after which microglia facilitate the extracellular conversion of ATP to adenosine, leading to activation of astrocytic A2B receptors and elevation of intracellular cAMP. This pathway provides a mechanistic explanation for the long-observed ability of antidepressants to enhance astrocytic cAMP/CREB signalling and supports growing evidence that astrocytes, microglia, purinergic signalling, and cAMP-dependent plasticity are central to antidepressant responses. More broadly, these data reinforce the view that SSRIs act not only through neuronal monoamine systems, but also by reshaping cooperative glial signalling networks that may underlie therapeutic recovery.

## Acknowledgements

The study was supported by the AMS Springboard award SBF005\1102 and the MRC Career Development Award MR/T031115/1 to VM.

We would like to thank Dr Stephen J Chuter for his assistance in generating the R scripts for analysis. We thank K. Jalink and A. Lann for providing the Epac-SH187 sensor clones, and O Griesbeck for donating the Twitch2B sensor clones.

## Code availability

The code used to calculate FRET ratio and normalised FRET ratio is available from the GitHub repository: https://github.com/marstoncatriona/FRETSensorAnalysis.

## Data availability statement

Data will be made available upon request to authors

## Funding statement

The financial support was provided by AMS Springboard award SBF005\1102 and the MRC Career Development Award MR/T031115/1 to VM

## Conflicts of interest

The authors report no conflicts of interest.

## Notes

### Competing Interest Statement

The authors have declared no competing interest.

