## Supplementary Tables for "Antidepressant fluoxetine engages astrocytic cAMP via purinergic signalling"

**Supplementary Table 1: Fluoxetine does not affect viability of primary rat astrocytes.**

| | Viable cells, % | | FLX vs Control |
|---|---|---|---|
| | Control | FLX | |
| 4 hours | 94.2±5.0 | 95.0±4.4 | t(3.93)=0.27; p=0.8460 |
| 24 hours | 99.0±0.8 | 98.1±0.9 | t(4.00)=1.34; p=0.2511 |

Primary rat astrocyte cultures were treated with fluoxetine (FLX; 10μM) for 4 or 24 hours. Cell viability was assessed using the trypan blue test. Data are shown as mean±SD; n=number of cells; N=number of wells; Control (4 hours) n=112; N=3; Control (24 hours): n=196, N=3; FLX (4 hours): n=92, N=3; FLX (24 hours): n=209, N=3; Unpaired t-test.

**Supplementary Table 2: Summary of target gene expression in primary rat astrocytes.**

| Gene | Expression, TPM |
|---|---|
| GFAP | 55810±11038 |
| Adora2b | 266±43 |
| Htr2b | 60±30 |
| Iba1 | 1994±241 |

RNAseq (BGI Genomics; paired-end, 100bp, 50 M clean reads per sample) has been performed on untreated primary astrocyte cultures. The table shows expression level of genes related to the current study: Glial Fibrillary Acidic Protein (GFAP; astrocytic marker); Adenosine 2B receptor (Adora2b); serotonin 2B receptor (Htr2b); and Ionised calcium-binding adaptor molecule 1 (Iba1; microglial marker). TPM, Transcripts Per Kilobase Million, a measurement of gene expression taking into account the gene length. Data are shown as mean±SD; n=5.

**Supplementary table 3: Serotonin 2B receptor expression in primary rat astrocytes.**

| Coverslip | Region | GFAP+/5-HT2BR+ cells, % |
|---|---|---|
| 1 | 1 | 100 |
| | 2 | 100 |
| | 3 | 100 |
| | 4 | 100 |
| | 5 | 100 |
| | 6 | 100 |
| 2 | 1 | 100 |
| | 2 | 100 |
| | 3 | 100 |
| | 4 | 100 |

Primary rat astrocytes were stained for Glial Fibrillary Acidic Protein (GFAP; astrocytic marker) and serotonin 2B receptor (5-HT2BR; BD Pharmigen 556334, 3μg/ml) as described in the main methods section "immunohistochemistry". Number of cells positive for both GFAP and 5-HT2B receptors were quantified from confocal images (Leica sp5; 20x air objective).
