## Supplementary Figure for "Antidepressant fluoxetine engages astrocytic cAMP via purinergic signalling"

**Supplementary Figure 1: Serotonin 2B receptor expression in primary rat astrocytes.**

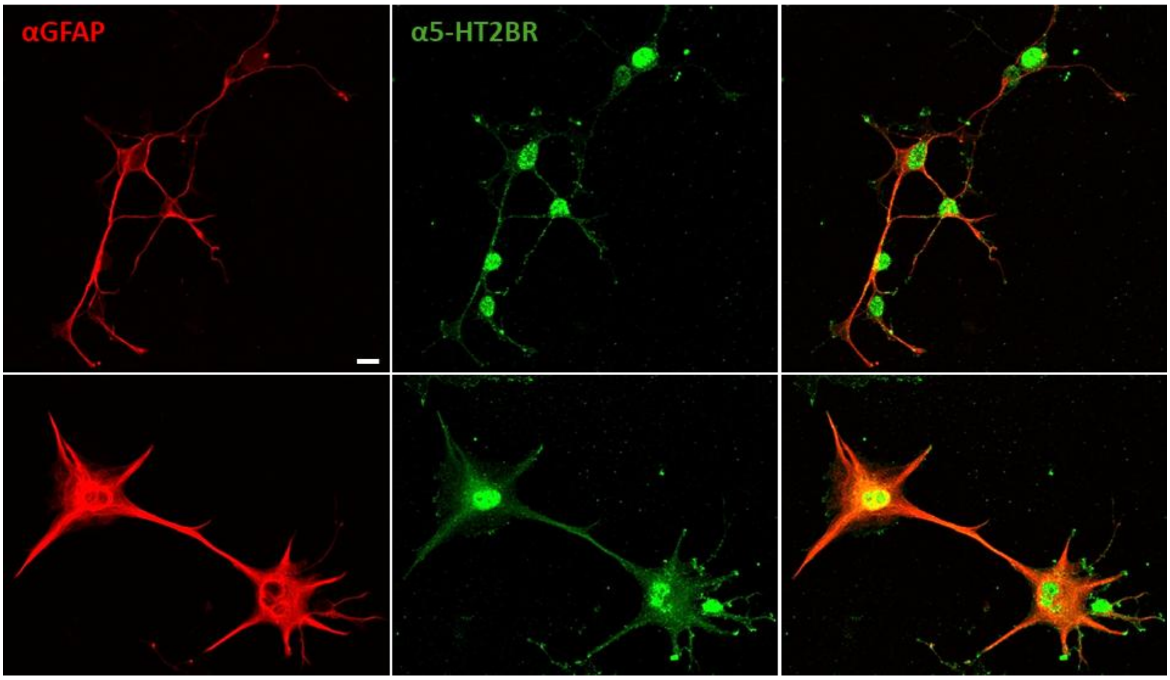


Representative images of primary rat astrocytes were stained for Glial Fibrillary Acidic Protein (GFAP; astrocytic marker) and serotonin 2B receptor (5-HT2BR). Scale bar = 10μm.
